# Potential depression and antidepressant-response biomarkers in human lymphoblast cell lines from treatment-responsive and treatment-resistant subjects: roles of SSRIs and omega-3 polyunsaturated fatty acids

**DOI:** 10.1101/2020.01.22.914358

**Authors:** Phatcharee Chukaew, Alex Leow, Witchuda Saengsawang, Mark M. Rasenick

**Author notes:** Correspondence: Department of Physiology and Biophysics, University of Illinois College of Medicine, 835 South Wolcott Avenue, Chicago, IL, 60612, USA.

## Abstract

While several therapeutic strategies exist for depression, most antidepressant drugs require several weeks before reaching full biochemical efficacy and remission is not entirely achieved in many patients. Therefore, biomarkers for depression and drug-response would help tailor treatment strategies. This study made use of banked human lymphoblast cell lines (LCLs) from normal and depressed subjects; the latter divided into remitters and non-remitters. Due to the fact that previous studies have shown effects on growth factors, cytokines and elements of the cAMP generating system as potential biomarkers for depression and antidepressant action, these were examined in LCLs. Initial gene and protein expression profiles for signaling cascades related to neuroendocrine and inflammatory functions differ among the three groups. Growth factor genes, including *VEGFA* and *BDNF* were significantly down-regulated in cells from depressed subjects. In addition, omega-3 polyunsaturated fatty acids (n-3 PUFAs) have been reported to act as both antidepressants and anti-inflammatories, but the mechanisms for these effects are not established. Here we showed that n-3 PUFAs and escitalopram (selective serotonin reuptake inhibitors, SSRIs) treatment increased adenylyl cyclase (AC) and *BDNF* gene expression in LCLs. These data are consistent with clinical observations showing that n-3 PUFA and SSRI have antidepressant affects, which may be additive. Contrary to observations made in neuronal and glial cells, n-3 PUFA treatment attenuated cAMP accumulation in LCLs. However, while lymphoblasts show paradoxical responses to neurons and glia, patient-derived lymphoblasts appear to carry potential depression biomarkers making them an important tool for studying precision medicine in depressive patients. Furthermore, these data validate usefulness of n-3 PUFAs in treatment for depression.

## Introduction

Depression is among the costliest and most disabling clinical maladies worldwide. The World Health Organization reported that about 4.4% of the world population, or over 300 million people suffer from depression [1]. Life time prevalence of depression is about one in six. Additionally, suicide represents a significant cause of death among those aged 15-59 years [2]. Despite this, the etiology of depression is still unclear, but it is known that both genetic and environmental factors play roles [3]. Although many treatment options are available, approximately one third of depressed subjects are classified as “treatment-resistant”, despite several drug switches or drug augmentations [4]. Also, even in patients that respond to traditional antidepressant therapy, an eight-week period of hysteresis exists between the onset of therapy and remission.

There is a critical need for peripheral biomarkers to be used in clinical applications for psychiatric disorders, especially for depression and antidepressant response. Growth factors and cytokines, as well as elements of cAMP signaling systems, represent highly promising candidates. Down-regulation of brain derived neurotropic factor (BDNF) [5–7] was observed, while vascular endothelial growth factor A (VEGFA), which can be both growth factor and inflammatory modulator, and interleukin 6 (IL6), which is a major pro-inflammatory cytokine, have been shown to be up-regulated in depression [8, 9]. Antidepressant treatments have been found to rectify these gene expression alterations [10, 11]. Furthermore, the system of molecules involved in the generation of cAMP might be potential biomarkers for depression and treatment response. The system regulating cAMP generation includes membrane-microdomain localization of the G protein, Gα and G-protein-coupled receptors (GPCRs). Additionally, cAMP generation is dependent upon the organization and accessibility of G proteins to receptors and effectors and other related membrane proteins such as caveolin-1 and flotilin-1 [12]. Localization of G alpha subunit (G_s_α) in lipid raft microdomains was attenuated in postmortem brain tissue from depressed subjects [13]. In addition, antidepressants were shown to translocate G_s_α from lipid raft to non-lipid raft [14], leading to robust coupling with its effector, adenylyl cyclase, resulting in increased cAMP production [15, 16].

Unfortunately, antidepressants can be accompanied by a series of side-effects that may be severe enough to cause patients to cease therapy. Therefore, it is important to discover additional compounds that may act as antidepressants, as well as adjuvants for conventional antidepressants to increase drug efficacy and decrease side-effects. Omega-3 polyunsaturated fatty acids (n-3 PUFAs) are commonly used to promote both somatic and mental health [17, 18]. Animals with PUFA-deficient diets showed the alteration of brain lipid compositions and depressed-like behaviors [19]. Peripheral n-3 PUFA levels in depressed subjects negatively correlate with symptom severity [20–22], suggesting that n-3 PUFAs might have beneficial effect for depressive patients. Consistent with these findings, clinical studies have been reported that PUFA mono-therapy is slightly beneficial over placebo [23]. The combination of selective serotonin reuptake inhibitors (SSRIs) and n-3 PUFAs augmented treatment efficacy in depression [24–26]. The exact mechanisms for n-3 PUFA effects on depression remain undetermined. The anti-inflammatory properties of n-3 PUFAs may be important for these antidepressant effects [27] since a high peripheral inflammatory index correlates with symptom improvement after n-3 PUFA administration [28]. Additionally, n-3 PUFAs that interact with membrane affect membrane “fluidity” and protein redistribution, resulting in alterations of downstream signaling cascades [29, 30]. Flotillin-1, a lipid-raft protein, was also displaced after n-3 PUFA treatment, suggesting that n-3 PUFAs can disrupt lipid rafts and modify cellular signaling.

In this study, we hypothesized that endogenous expressions of growth factor, inflammatory cytokine and cAMP generating component can be used as biomarkers in depression and antidepressant response. In addition, those targets might underlie the antidepressant mechanism. We used human lymphoblast cell lines (LCLs) derived from healthy controls and depressed subjects as cell models. LCLs are used for precision medicine studies of therapeutics in brain disorders, including depression [31–33]. Thus, we investigate depression and drug-response biomarkers in LCLs, comparing healthy controls, depression subjects who remitted to antidepressant treatment and those who did not remit. We also sought to use this platform to examine antidepressant-like action of n-3 PUFAs, with and without SSRI, in modifying signaling cascades and membrane protein distribution. We found that expression profiles of depression risk genes and proteins in LCLs from depressed subjects differed from those in healthy controls. Some of those genes can be potential biomarkers for depression and antidepressant response. Furthermore, n-3 PUFA alone and combination with escitalopram affected membrane microdomain modification and some related signaling cascades that might underlie antidepressant-like action of n-3 PUFAs for treatment in depression.

## Materials and methods

### 1. Cell culture

Human lymphoblast cell lines (LCLs) from healthy control and depressed subjects were obtained from the NIMH. LCLs from healthy controls (n=10) were obtained from study 76: Genomic Psychiatry Cohort (GPC) while LCLs from depressed subjects (n=24) were obtained from study 18: Sequenced Treatment Alternatives to Relieve Depression (STAR*D). LCLs derived from patients used in this study were from mild to moderate depression with Hamilton rating scale (HRS) for depression ranging from 14 to 18 scores during enrollment. HRS of each patients after treatment was used to divide group of pretreated LCLs that from depressed subjects who remitted (HRS score after treatment ≤ 7, n=12) and did not remit (HRS score after treatment > 7, n=12) to antidepressant treatment (see supplementary table 2 and 3 for complete description). Cells were maintained under humidified air containing 5% CO_2_ at 37°C in incubator. LCLs were grown in RPMI-1640 with 10% vol/vol of fetal bovine serum and 1% vol/vol penicillin-streptomycin. Cells were plated into T25 flask at 1 x 10^5^ cells/ml in cultured media. One day following plating, cells were treated with vehicle, 20 μM PUFAs (1:1 of EPA and DHA (U-99-A and U-84-A, respectively from NU-CHECK-PREP)), 10 μM escitalopram (LU 26-054-0, Drugbank) or 20 μM PUFAs + 10 μM escitalopram for 72 hours. In other sets of treated conditions, cells were plated into T25 flasks at 4 x 10^5^ cells/ml. One day following plating, cells were treated with vehicle, 10 or 20 mM methyl-β-cyclodextrin (MBCD) (C4555, Sigma) for 30 minutes or vehicle, 2 or 10 μM ketamine for 60 minutes. Cell density on the day of collection was approximately at 8 x 10^5^ cells/ml. Cells were transferred to 50 ml tubes, centrifuged at 2000xg for 2 minutes at room temperature, and washed twice with PBS before performing RNA extraction, total protein extraction, membrane fractionation or cAMP assay.

### 2. Target selection: Genes and proteins

All genes and gene products investigated were selected prior to initiation of experiments. Previous studies have focused on lipid rafts (hence caveolin and flotillin). cAMP generation (hence G_s_α, G_olf_α and isoforms of adenylyl cyclase), cAMP-regulated growth factors (hence VEGFA and BDNF) and inflammatory mediators linked to n-3 PUFA (hence IL6). Taken together, these markers were considered a broad representation of our target.

### 3. RNA extraction and qRT-PCR

RNA was extracted using Trizol reagent (15596018, Ambion by Life Technologies). Briefly, cells were collected and the pellets were lysed with 1 mL of Trizol reagent. The insoluble components were removed by centrifugation at 12,000xg for 10 minutes at 4°C. The supernatant was transferred into a new microcentrifuge tube and incubated for 5 minutes at room temperature. To precipitate DNA, 200 µL of chloroform was added into the tube. Then, the tube was shaken vigorously by flipping the tube for 15 seconds and incubating for 3 minutes at room temperature. The samples were centrifuged at 12,000xg for 5 minutes at 4°C to separate into three phases. The aqueous phase (top layer) containing RNA was transferred into a new microcentrifuge tube and RNA was precipitated by adding 500 µL of isopropanol, incubation for 10 minutes at room temperature followed by centrifugation at 12,000 rpm for 10 minutes at 4°C. The supernatants were discarded. The RNA pellet was washed with 1 mL of 75% ethanol and dried at room temperature. To solubilize RNA, RNase free water was added. The RNA concentration was measured using a Spectramax i3X microplate reader and the SolfMax Pro software (Molecular Devices) to make 1 mg/ml RNA concentration before storage at -80°C.

cDNA was synthesized from total RNA isolated from LCLs by using a reverse transcription kit iScript^TM^ cDNA synthesis (1708891, Biorad). The reaction was performed according to the manufacturer’s instructions. The reaction without reverse transcriptase enzyme was used as a negative control. Finally, cDNAs were stored at -20°C until used for quantitative real-time polymerase chain reaction (qRT-PCR). For cDNA amplification, cDNA was mixed with PowerUp SYBR Green Mastermix reagent (4367659, ThermoFisher), forward and reverse primers (as shown in Supplementary table 1), and nuclease-free water. Samples were analyzed in triplicate and run on Life Technologies ABI ViiA7 RT-PCR equipment with Quantstudio real time PCR software. The temperature set up was started with the holding stage at 95°C for 10 minutes, followed by 40 cycles of amplification by 95°C for 15 seconds and 60°C for 1 minute, and created melting curve by 95°C for 15 seconds, 60°C for 1 minute and 95°C for 15 second. Values of sample containing the target genes were calculated by the formula E = 10^(−1/slope)^, E = 2 when efficiency correction is 100%. The expression ratio of target gene were determined by the mathematical model as 2^-ΔCt^ when ΔCt = Ct target gene - Ct reference gene. The relative expression ratio to control of target gene from the treatment effect was determined by 2^-ΔΔCt^ when ΔΔCt = ΔCt control - ΔCt treatment.

**Table 1.** Human lymphoblast cell lines from healthy control and depressed subjects who remitted and or did not remit to antidepressant treatment

| Group | Gender (n) | Age (years) | HRS |  |
| --- | --- | --- | --- | --- |
|  |  |  | Before treatment | After treatment |
| Control | Male n = 5 | 40.60 ± 5.44 | - | - |
|  | Female n = 5 | 32.60 ± 5.83 | - | - |
|  | Total n=10 | 36.60 ± 3.99 | - | - |
| Remitter | Male n = 5 | 39.60 ± 6.78 | 16.40 ± 0.60 | 3.80 ± 0.97 |
|  | Female n = 7 | 40.57 ± 3.80 | 16.71 ± 0.29 | 3.86 ± 0.91 |
|  | Total n=12 | 40.17 ± 3.40 | 16.58 ± 0.29 | 3.83 ± 0.64 |
| Non- remitter | Male n = 6 | 44.67 ± 4.12 | 16.50 ± 0.67 | 13.50 ± 1.93 |
|  | Female n = 6 | 50.67 ± 4.11 | 16.67 ± 0.49 | 17.00 ± 3.44 |
|  | Total n=12 | 47.67 ± 2.92 | 16.58 ± 0.40 | 15.25 ± 1.95 |

Means ± SEM are provided where applicable, – = not applicable

### 4. Cell lysate and Western blotting

For basal protein expression determination, cells were lysed with 1XRIPA lysis buffer (9806S, Cell signaling) containing 1X proteinase inhibitor (PI) cocktail (P2714-1BTL, Sigma). Lysed cells were homogenized and incubated on ice for 15 minutes. The insoluble components were removed by centrifugation at 12,000xg for 15 minutes at 4°C. To separate lipid raft and non-raft membrane fractions, fractionation using Triton X-100 (28314, Fisher) and Triton X-114 (28332, Fisher) was used (modified from Toki et. al, 1999 [34]). Treated cell pellets were re-suspended with TME buffer containing 1X PI. The cell solution was homogenized and incubated for 30 minutes on ice. Part of the resulting solution was collected as the whole cell lysate fraction, and the rest was centrifuged at 57,300xg for 60 minutes at 4°C. The supernatant was collected into a microcentrifuge tube labeled as cytosolic protein. Next, pellet was re-suspended with 1% Triton X-100 in TME with 1X PI, homogenized and recentrifuged as mentioned above. The supernatant was collected into a microcentrifuge tube labeled as TX-100 soluble membrane or non-lipid raft fraction. The pellet was re-suspended again with 1% Triton X-114 in TME with 1X PI before homogenization and centrifugation as stated above. Note that this extraction procedure gives results similar to “detergent-free” extraction methods (high pH treatment can result in some saponification) or sucrose-density gradients [35]. The supernatant was collected into a microcentrifuge tube labeled as TX-114 soluble membrane or lipid raft fraction. The final pellet was discarded. A BSA assay kit was used to measure protein concentration of samples before performing western blotting analysis by using the Protein simple Wes system. The EZ Standard pack 1 (PS-ST01EZ-8) was used to mix with 4 μg total protein. The 12-230 kDa plate and 8×25 capillary cartridge (SM-W004-1), anti-Mouse Detection module (DM-002) and anti-Rabbit Detection module (DM-001) from Protein Simple were used with specific primary antibodies, including 1:400 anti-G_s_α protein (75-211, Neuromab), 1:100 anti-GNAL (G_olf_α protein (PA5-50981, Invitrogen), 1:100 anti-Flotilin-1 (3253, CST), 1:400 anti-beta actin (3700, CST), 1: 100 anti-VEGFA (ab46154, abcam) and 1:40 anti-proBDNF (sc65514, Santa cruz). The plate was run with the Protein Simple WES machine and data analyzed with Compass software.

### 5. Alphascreen cAMP

Alphascreen cAMP Assay kit (6760635M, PerkinElmer) was used to measure cAMP in the samples. Treated cell pellets were re-suspended in stimulation buffer that was prepared fresh from 1x HBSS, 5 mM HEPES, 0.5 mM IBMX and 0.1% BSA. cAMP assays were performed following manufacturer’s instructions and forskolin (10 μM final concentration) was added into each well to stimulate cAMP production. A cAMP standard curve was created from fluorescent signals of serial dilutions of cAMP standard solution. A Spectramax i3X microplate reader with alphascreen module and SolfMax Pro software (Molecular Devices) was used for signal detection and to measure cAMP accumulation.

### 6. Statistical analysis

Outlier(s) were identified by ROUT (Q = 1%) before testing the distribution or variances by D’Agostino & Pearson omnibus normality test. If variances were equal or data were normally distributed, one-way ANOVA followed by Bonferrroni’s multiple comparisons test were performed to compare treatment effects (3 pair-wise comparisons; Healthy vs. Remitters, Healthy vs. Non-remitters, and Remitters vs. Non-remitters). One-way Kruskal–Wallis test followed by Dunn’s multiple comparisons test was performed for unequal variances or non- nonnormal distributed data. Results are represented as mean ± SEM. Statistical analysis was performed with the GraphPad Prism version 6.0 software (GraphPad Software Inc., San Diego, CA).

## Results

### 1. Human lymphoblast cell lines (LCLs) reveal potential depression biomarkers

Human LCLs have been used for studying a variety of neuropsychological diseases including depression [32, 33]. To determine whether LCLs possess depression and drug-response biomarkers, we compared the expression of signaling genes and proteins, previously determined to be relevant to depression and antidepressant response, in LCLs from control and depressed subjects [5-8, 10-14, 36]. Pre-treated LCLs were developed from peripheral blood of subjects during the enrollment step for each study. Cells from depressed subjects underwent antidepressant treatment were divided into 2 groups based on results from the Hamilton rating scale for depression (HRS), which includes 17 items. Cells were grouped into those from subjects who remitted after sequenced treatment (HRS ≤7; remitters) and those who still had depressed symptoms (HRS >7; non-remitters). Cell lines from all groups were isolated from both male and females aged between 22 to 63 years. Prior to treatment, patients HRS scores were not significantly different between sexes and treatment response (see Table 1, for complete information on subjects from whom LCLs were obtained, see Supplementary table 2 and 3).

Previous evidence has shown that both flotilin-1 and caveolin-1 are enriched in lipid raft microdomains, and used as lipid raft markers, during antidepressant treatment [37]. Expression levels of flotillin-1 protein (Supplementary figure 1a and 1b), but not *FLOT1* mRNA (Fig. 1a) were significantly lower in LCLs from depressed subjects compared with healthy controls. Interestingly, contrary to our results in LCLs, a previous study showed up-regulation of *FLOT1* in the brain of depressed patients [38]. Differences between *FLOT1* (gene) and flotilin-1 (protein) may reflect posttranslational modification or targeted degradation in depressed subjects. Caveolin-1 protein was undetectable in LCLs. However, we did detect *CAV1* mRNA, and observed a down-regulation of *CAV1* in LCLs from remitters (Fig. 1b).

**Fig. 1.**
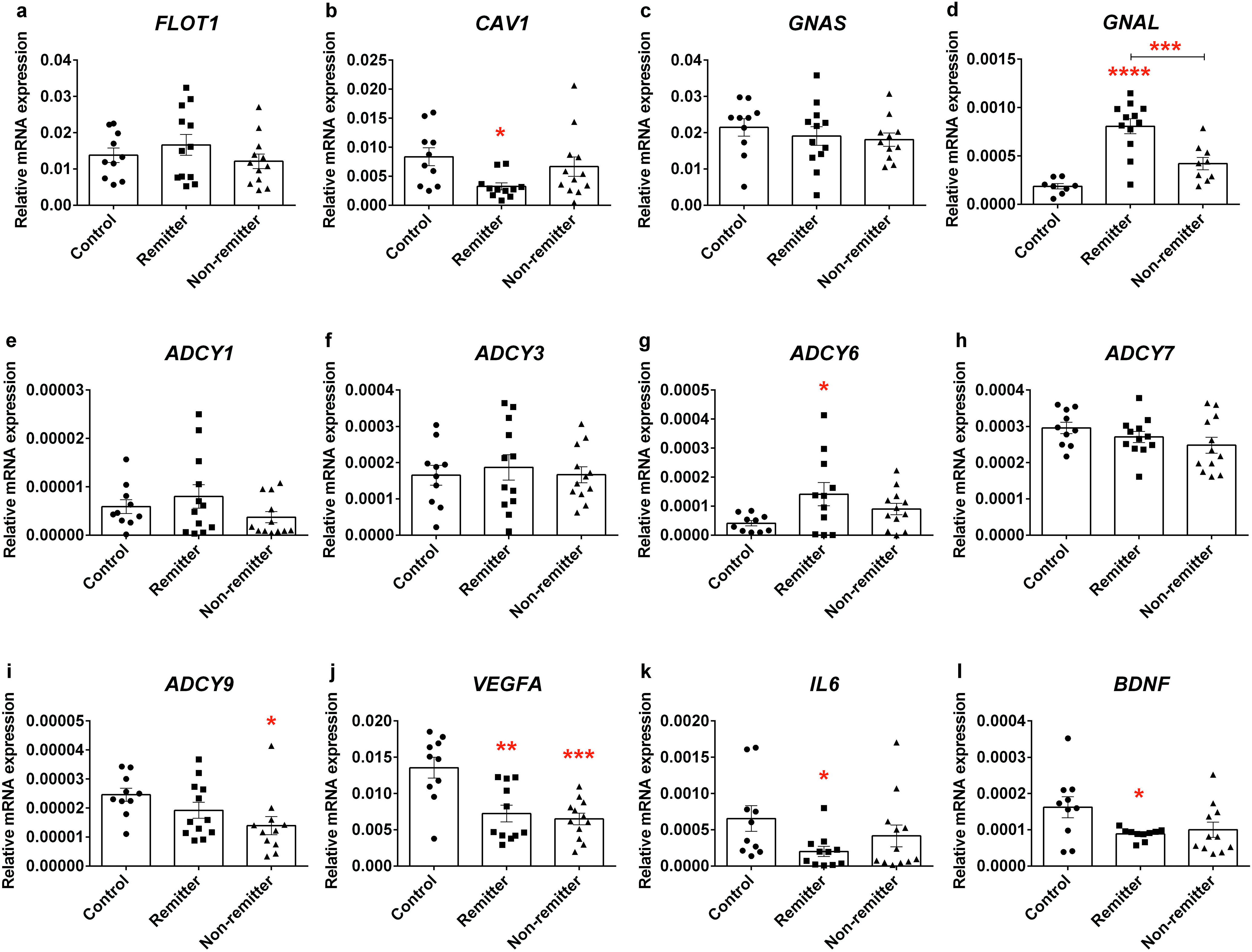
Basal expression levels of genes in LCLs from healthy controls and depressed subjects who responded and did not respond to antidepressant treatment. Total RNA was collected and used for reverse transcription. qRT-PCR was used to measure the target gene expression including **a** *FLOT1*, **b** *CAV1*, **c** *GNAS*, **d** *GNAL*, **e** *ADCY1*, **f** *ADCY3*, **g** *ADCY6*, **h** *ADCY7*, **i** *ADCY9*, **j** *VEGFA*, **k** *IL6* and **l** *BDNF*. Each experiment was done in triplicate and expression levels were normalized with beta actin (2^-ΔCt^). All genes and gene products investigated were selected a priori as previous studies have focused on lipid rafts (hence Caveolin and flotillin), cAMP generation (hence Gs, Golf and isoforms of adenylyl cyclase), cAMP/regulated growth factors (hence VEGFA and BDNF), and inflammatory mediators linked to n-3 PUFA (hence IL6). Values from each cell line are represented as dots in the scatter plot-bar of mean ± SEM. Adjusted *p* value from 3 post-hoc pair-wise comparisons among groups of cell lines, \**p*<0.05, \*\**p*<0.01, \*\*\**p*<0.001, \*\*\*\**p*<0.0001.

The cAMP generating system has long been invoked as relevant to depression. Congruently, a recent PET study found that lowered cAMP in depressive patients is rectified subsequent to effective antidepressant treatment [39]. We have suggested that G_s_α signaling, a known mediator for cAMP production, is attenuated in depression [13] due to excessive lipid raft localization, which is reversed in response to antidepressants [16, 35] without changes in the expression of G_s_α (or other Gα subunits) [34, 40] in brain, neurons or glia. In the current study, we measured mRNA and protein expression of G_s_α and G_olf_α the stimulatory G protein family, in the LCLs. While *GNAS*, which encodes G_s_α, was not altered in cells from the depressed subjects, *GNAL* which encodes G_olf_α, were higher in the LCLs from remitter (Fig. 1c and 1d) relative to control. However, there was no significant difference for protein expression among cell lines (Supplementary figure 1a, 1c and 1d).

Five adenylyl cyclase (AC) isoforms, which catalyze the production of cAMP from ATP, genes could be detected in human LCLs, including *ADCY1*, *ADCY3*, *ADCY6*, *ADCY*7, and *ADCY9* (Fig. 1e-1i). Among these genes, *ADCY7* and *ADCY3* were highly abundant in LCLs from all groups but were not significantly different in depressed patients compared to control (Fig. 1f and 1h). While *ADCY1* was not altered in LCLs from depressed subjects (Fig. 1e), *ADCY6* was up-regulated in LCLs from remitters relative to those from healthy controls (Fig. 1g). In contrast, *ADCY9* expression was lower in LCLs from non-remitters compared to healthy controls (Fig. 1i). Taken together these findings suggested that *GNAL* and some ADCY isoforms might be potential antidepressant-response biomarkers.

BDNF is relevant for both depression and antidepressant response [41]. Decreased *BDNF* mRNA expression was detected in LCLs from remitters (Fig. 1l). There was no difference in proBDNF, a BNDF precursor, expression among cell lines (Supplementary figure 1a and 1f). However, BDNF ELISA showed a significantly up-regulation of total BDNF protein in LCLs derived from depressed subjects (Supplementary figure 1g). Differences observed in *BDNF* gene expression and total BDNF protein in LCLs from depressed subjects illustrate its potential as a peripheral biomarker for depression, as well as for examining the efficacy of antidepressant treatment.

Dysregulation of immune function has been proposed to be relevant to the etiology of depression [42, 43]. Therefore, we determined the expression of VEGF and IL6, which are categorized as major cytokines that have been reported to be altered in depressed patients [8–11]. We found that *VEGFA* gene (Fig. 1j) but not protein (Supplementary figure 1a and 1c) was significantly down-regulated in LCLs from depressed subjects. We also found down-regulation of *IL6* mRNA in LCLs from remitters (Fig. 1k). IL6 protein expression was also undetectable. Both VEGFA and IL6 have been considered as potential risk genes for depression [44]. Most studies report up-regulation of both *VEGFA* and *IL6* in the depressive brain [8, 9], we discovered the opposite in LCLs. However, this was also seen in serum and CSF of depresses patients [45, 46].

### 2. PUFAs and escitalopram altered gene expression in LCLs derived from depressed subjects

Previous studies have reported that PUFAs may improve depressive symptoms. Omega-3 PUFA can augment the efficacy of SSRI and might have intrinsic anti-depressant properties [23, 24]. To determine whether n-3 PUFAs with and without escitalopram, an SSRI, affect expression levels of genes related to depression, the LCLs from both the remitters (n=5) and non-remitters (n=4) were pooled and treated with vehicle, 20 μM PUFAs, 10 μM escitalopram or 20 μM PUFAs + 10 μM escitalopram for 72 hours before RNA isolation. *FLOT1* and *BDNF* expressions were increased by n-3 PUFAs and escitalopram combination (Fig. 2a and 2l). Neither n-3 PUFAs nor escitalopram treatment had significant effect on *CAV1*, *GNAS*, *GNAL*, *ADCY6* and *ADCY9* genes (Fig. 2b-d, 2g and 2i). Omega-3 PUFAs alone significantly increased *ADCY7*, (Fig. 2h) while escitalopram alone increased *ADCY1* and *ADCY3* expressions (Fig. 2e and 2f).

**Fig. 2.**
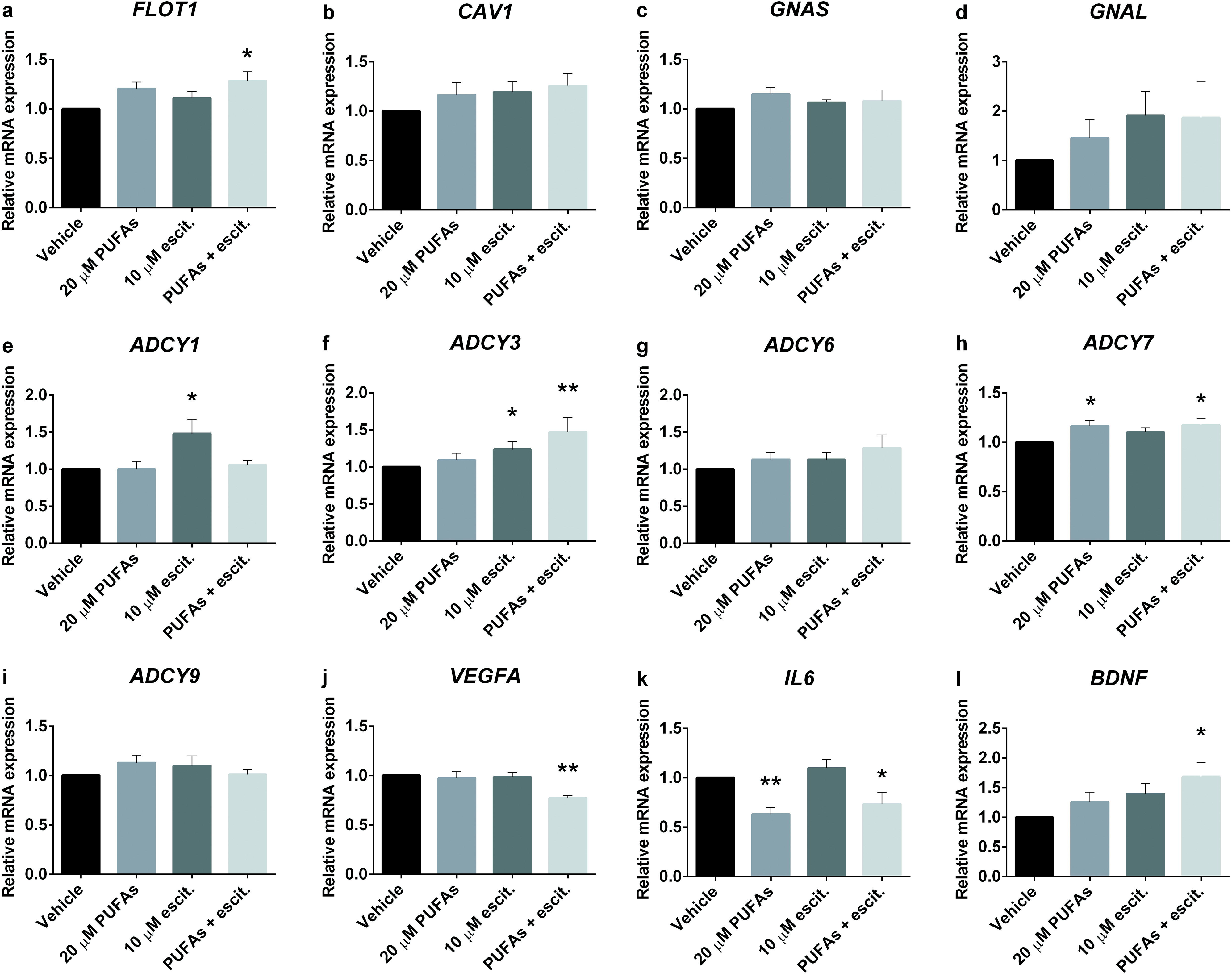
PUFA and escitalopram treatment-altered gene expression in LCLs derived from depressed subjects. Total RNA was collected from LCLs that were treated with vehicle, 20 μM PUFAs, 10 μM escitalopram or 20 μM PUFAs + 10 μM escitalopram for 72 hours. qRT-PCR was used to measure target gene expression including **a** *FLOT1*, **b** *CAV1*, **c** *GNAS*, **d** *GNAL*, **e** *ADCY1*, **f** *ADCY3*, **g** *ADCY6*, **h** *ADCY7*, **i** *ADCY9*, **j** *VEGFA*, **k** *IL6* and **l** *BDNF*. The experiment was carried out in 10 cell lines. Values of relative expression to control (2^-ΔΔCt^) indicate bar of mean ± SEM. Adjusted *p* value of 3 post-hoc pairwise comparisons between treated conditions and vehicle, \**p*<0.05, \*\**p*<0.01.

The effects of n-3 PUFAs and escitalopram on cytokine mRNA expression were also determined. The *VEGFA* expression was significantly decreased in the n-3 PUFAs and escitalopram combination (Fig. 2j). In addition, *IL6* expression was suppressed by n-3 PUFAs (Fig. 2k). Taken together these data suggested that n-3 PUFAs, in combination with escitalopram might affect membrane structure, cAMP generating cascade, growth factor and pro-inflammatory cytokine expression. These data lend a molecular rationale to SSRI + n-3 PUFA therapeutics for depression.

### 3. PUFAs and escitalopram altered protein expression and membrane distribution in LCLs derived from depressed subjects

A consistent antidepressant biosignature is the translocation of G_s_α protein from lipid rafts to non-raft membrane domains, where it more effectively activates adenylyl cyclase [14]. To test whether the “antidepressant action” of n-3 PUFAs includes the redistribution of membrane proteins, LCLs from depressed subjects were treated as described above and then subjected to biochemical fractionation and western blotting. While the treatments did not change total expression levels of neither flotillin-1 (Fig. 3a-b) nor G_olf_α, (Fig. 3i and j) in the LCLs, both n-3 PUFAs and escitalopram increased flotillin-1 in non-lipid raft (Fig. 3a and 3c), but not lipid raft microdomains (Fig. 3a and 3d), suggesting that the treatment might alter the lipid raft structure and component. In addition, n-3 PUFAs alone increased G_olf_α on both non-lipid raft and lipid raft areas (Fig. 3i, 3k and 3l).

**Fig. 3.**
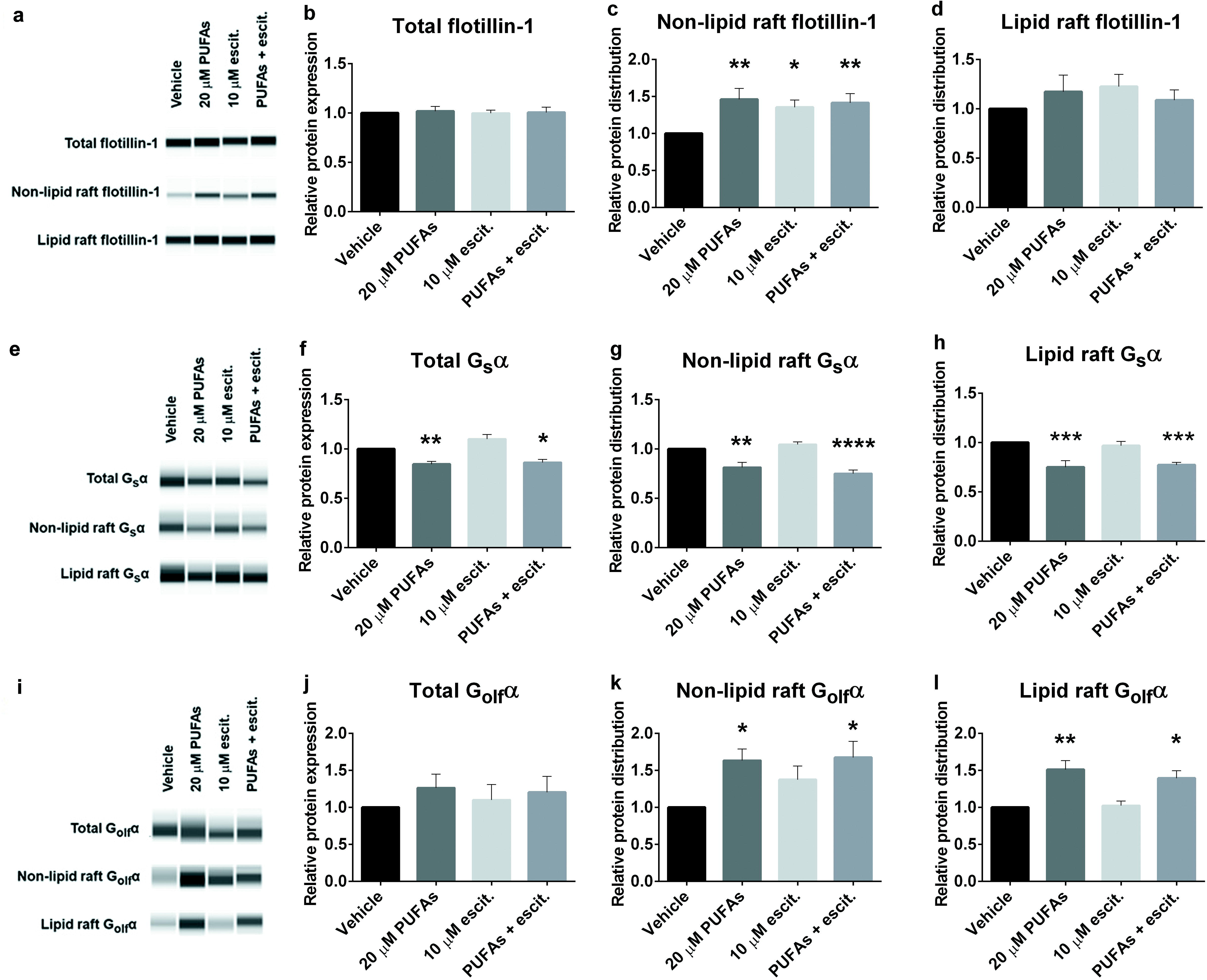
PUFAs and escitalopram altered G protein and flotillin distribution in LCLs derived from depressed subjects. Biochemical fractionation using TX100/TX114 was performed to isolate whole cell lysate, non-lipid raft and lipid raft fractions from LCLs that were treated for 72 hours with vehicle, 20 μM PUFAs, 10 μM escitalopram or 20 μM PUFAs + 10 μM escitalopram. Protein concentrations of each sample were equalized before loading the same amount into a Protein Simple Wes plate for capillary electrophoresis/western blotting. The experiment was done in 9 cell lines. Sample Wes results are shown in a,e and i. Values for mean ± SEM of **a**-**d** Flotillin-1, **e**-**h** G_s_α and **i**-**l** G_olf_α. Adjusted *p* value of 3 post-hoc pairwise comparisons between treated conditions compared with vehicle, \**p*<0.05, \*\**p*<0.01, \*\*\**p*<0.001, \*\*\*\**p*<0.0001.

In contrast, n-3 PUFAs decreased total G_s_α expression (Fig. 3e and 3f), leading to decreased G_s_α in both non-lipid raft and lipid raft domains (Fig. 3e, 3g and 3h). Decreased total G_s_α expression and membrane distribution were also observed in LCLs treated with the lipid raft disruption agent, MBCD (Supplementary figure 3a-d). These findings supported the hypothesis that n-3-PUFAs redistribute membrane proteins as an underlying mechanism of antidepressant-like action. However, the redistribution of membrane proteins is different in LCLs relative to neuronal or glial cell populations. This is not observed after treatment with other antidepressants, including ketamine (Supplementary figure 3e-3h), in contrast with studies on C6 cells and primary astrocytes [34, 47], suggesting the cell type-specific effects of antidepressant treatment.

Chronic SSRIs treatment changed peripheral protein levels such as BDNF [48] and VEGFA [11]. Then, we further examined roles of n-3 PUFAs and escitalopram in cytosolic VEGFA and proBDNF levels in LCLs from depressed patients. Treated conditions have no effect on VEGF and proBDNF expression (Supplementary figure 2a-2c).

### 4. PUFAs and escitalopram decreased cAMP accumulation in LCLs

Distribution of membrane proteins especially, G_s_α, affects cellular cAMP accumulation [15]. We found that n-3 PUFAs alone decreased G_s_α membranes association. Also, n-3 PUFAs co-treatment with escitalopram increased non-lipid raft flotillin-1, a lipid raft protein. Next, to investigate whether n-3 PUFAs and escitalopram co-treatment alters cAMP accumulation, we determined cAMP levels in treated LCLs. Forskolin stimulation, which indicates G_s_α/AC association [49], significantly increased cAMP accumulation from 0.32±0.03 to 1.85±0.35 nM per 10,000 cells of each cell line (n=20). However, this did not vary significantly among cells in each group (Fig. 4a). Accumulation of cAMP after treatment was normalized to the basal level of each cell line and represented as relative cAMP level in the presence of forskolin stimulation. Omega-3 PUFAs or escitalopram treatment alone significantly decreased relative cAMP accumulation in LCLs, but no additive effects were observed (Fig. 4b). Given the paradoxical effects of these compounds showed on cAMP accumulation in LCLs relative to neurons and glia [50, 51], we further tested effects of MBCD and ketamine on cAMP accumulation [15, 47]. We found that the MBCD-induced translocation of G_s_α from membrane domians reduced cAMP accumulation in LCLs while ketamine treatment had no effect on G_s_α translocation of cAMP accumulation (Supplementary figure 4a and 4b).

**Fig. 4.**
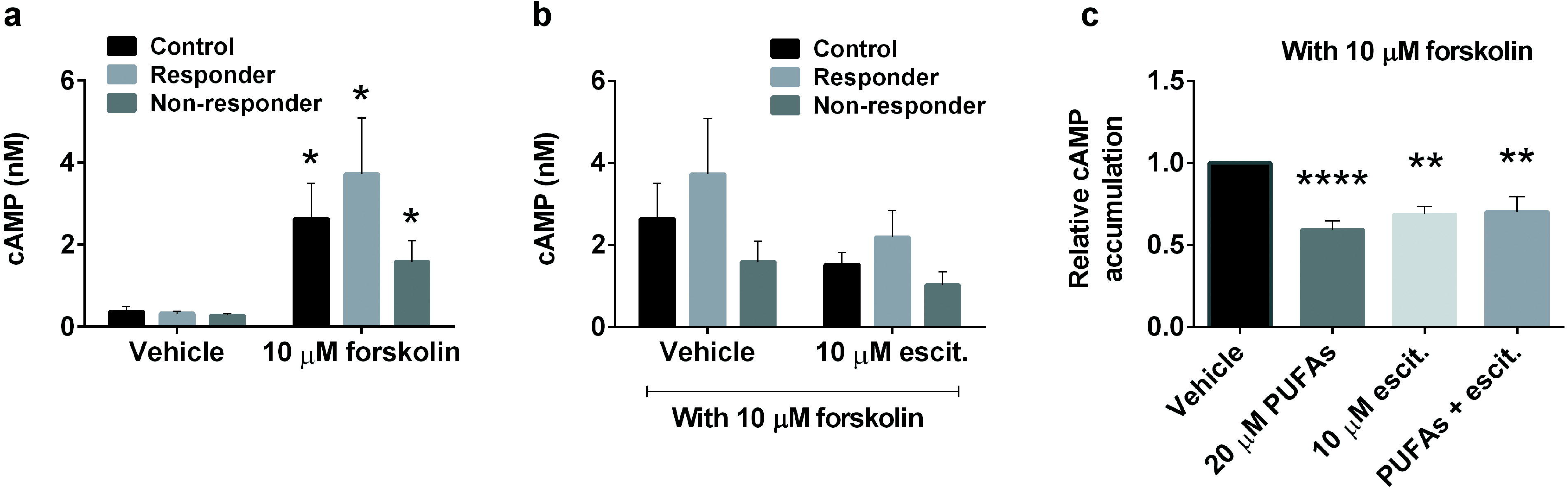
PUFAs and escitalopram decreased cAMP accumulation in LCLs. Alpha screen cAMP was used to measure cAMP accumulation in cells, and values are derived from cAMP standard curves developed in each assay. LCLs that were treated for 72 hours with vehicle, 20 μM PUFAs, 10 μM escitalopram or 20 μM PUFAs + 10 μM escitalopram were collected and used for cAMP assays. **a** Forskolin significantly increased cAMP accumulation in LCLs from all groups, including, healthy control, remitters and non- remitters. **b** Post-treatment cAMP accumulation with forskolin was compared to vehicle control and was done in triplicate for 20 cell lines derived from all group of cell lines. Values indicate mean ± SEM. **a** two-tail *p* value and **b** adjusted *p* value of post-hoc 3 pairwise comparisons between treated conditions compared with vehicle, \**p*<0.05, \*\**p*<0.01, \*\*\*\**p*<0.0001.

To our surprise, LCLs displayed both gene expression profiles and antidepressant- treatment responses opposite (for the most part) to results previously observed in neurons and glia. SSRI co-treatment with n-3 PUFAs caused alterations in membrane distribution of flotillin- 1, G_s_α and G_olf_α proteins, resulting in attenuated cAMP accumulation in human LCLs. Direct disruption of lipid rafts with a cholesterol chelator had opposing effects in LCLs (decreased coupling between G_s_α and AC) as in neurons or glia. This result was consistent with the opposing effects of antidepressants on cAMP signaling parameters observed in LCLs. Taken together these data suggest that patient-derived lymphoblasts carry potential depression biomarkers and might be a good model for studying individual drug responses and personalized medicine. Furthermore, these data validate the usefulness of n-3 PUFAs in treatment for depression and propose possible mechanisms underlying antidepressant-like effect of n-3 PUFAs.

## Discussion

This project was undertaken to determine the usefulness of LCLs as cellular models for depression and antidepressant response. We also sought to determine if this system had potential to study the antidepressant effects of n-3 PUFAs, either independently or in combination with SSRIs. Human LCLs from patients with depression showed differential expression of potential depression risk genes in comparison with LCLs from healthy controls. These data suggest that LCLs might be useful probes to help design a personalized therapy plan for depressive individuals.

Expression of specific genes and proteins related to cAMP-signaling and cAMP generating genes in LCLs from depressed subjects differed from those in healthy controls (Fig. 1). We found flotillin-1 protein expression levels were significantly lower in LCLs from depressed subjects compared with healthy controls. However, *FLOT1* mRNA expression did not differ in LCLs from depressed subjects versus healthy controls (compare Fig. 1a and supplementary figure 1a and 1b). However, it is important to note that mRNA and protein levels are not always directly correlated, particularly among proteins relevant to G protein signaling [52]. Furthermore, protein degradation might be altered because of the deregulation of ubiquitin-proteasome system in depression [53]. Flotillin-1 can interact with the serotonin transporter (SERT). Deficiencies in flotillin-1 lead to an increased SERT activity, thereby altering serotoninergic signaling [54]. In addition, flotillin-1 is important for inflammatory signaling [55]. Expression of flotillin-1 protein was suppressed in LCLs from depressive subjects, suggesting its use as a potential peripheral marker for depression. Curiously, *FLOT1* was significantly up- regulated in brains and peripheral blood of depressive cases compared with controls [38]. This was one of the clear differences we noted between LCLs and neuronal and glial cells.

Caveolin-1 is located mainly in lipid raft microdomains in many different cell types [56]. Recent studies revealed that caveolae and caveolin are present in all immune cells depending on their activation state [57]. Caevolin-1 was undetectable in LCLs, which are non-activated immune cells. However, expression of *CAV1* mRNA was clearly detected and significantly down-regulated in LCLs from remitters (Fig. 1b). Expression and localization of signaling components on plasma membrane microdomains are relevant to regulation of cellular signaling pathways [58]. Thus, alteration of *CAV1* expression might regulate immune response in a manner relevant to depression.

Dysfunction of the cAMP generating systems has been found in depression [59] and is resolved by successful antidepressant treatment [39]. We found that expression of *GNAL* and some *ADCY* genes differed in LCLs from depression subjects versus controls (Fig. 1). Adenylyl cyclases generate cAMP and are activated by G_s_α or G_olf_α, the latter primarily for AC3 [60]. Adenylyl cyclase can be quantified through mRNA or activity (which does not give information on specific isoforms), but antibodies are both non-specific and non-quantitative in this study. Although expression levels of *GNAL* and some *ADCY*s were not uniform among groups of LCLs, basal cAMP levels and forskolin-activated adenylyl cyclase were not different (Fig. 4a). AC7 plays critical roles immune cells [61] and showed the most robust expression in LCLs (Fig. 1h). AC7 is not abundant in neurons or glia, and is absent from C6 cells. *ADCY7* expression in LCLs was not significantly different among individual lines, and this may underlie the consistencies in basal and forskolin-activated cAMP accumulation (Fig. 4a). A previous study also reported that cAMP levels in leucocytes from healthy control and depressive subjects were not different [62], which is consistent with our findings (fig. 4a). However, the same study showed less cAMP accumulation by GPCR agonist stimulation in leucocytes from depressed subjects. This difference remains unresolved, but current methods allow cAMP determination with greater accuracy and precision.

Immune dysfunction has been suggested to be correlate with depression [63]. To further support this hypothesis, we showed down-regulation of *VEGFA* and *IL6* expression in LCLs from depressed subjects. VEGFA which is an important angiogenesis factor has also been categorized as an inflammatory cytokine. VEGFA can be both pro-inflammatory and anti-inflammatory [64]. Unlike VEGFA, IL6 plays a major role as a pro-inflammatory cytokine, which has also been implicated in depression [65]. The majority of studies have reported an up- regulation of central and peripheral *VEGFA* and *IL6* in depression [8, 9, 64, 65]. Conversely, we found lower *VEGFA* and *IL6* in LCLs from depressed subjects versus those from healthy controls (Fig. 1j and 1k). Possibilities for this discrepancy may due to the heterogeneity of depression, as well as the differences we noted between LCLs and neuronal and glial cells. As we have shown that multiple immune system targets were altered in cells from depressed subjects, *VEGFA* and *IL6* might be depression risk genes. As we hypothesized, antidepressant-like effect of n-3 PUFAs might have an anti-inflammatory component. We found that n-3 PUFAs suppressed *IL6* expression (Fig. 2k) and the combination of n-3 PUFA and escitalopram suppressed *VEGFA* expression (Fig. 2j). These findings were consistent with a previous study reporting that n-3 PUFAs down-regulated expression of cytokines, including *VEGFA* and *IL6* [66].

BDNF expression levels have been implicated in regulating depression and antidepressant action [41]. Expression of peripheral BDNF correlates with CNS BDNF expression, therefore, alteration of blood BDNF might reflect similar action in the CNS [67]. Several antidepressants, including escitalopram, can increase BDNF [35]. Alterations in BDNF have potential to use as a prognostic biomarker [68]. Consistent with these studies, we found a down-regulation of *BDNF* in remitters (Fig. 1l). However, total BDNF, measured by ELISA, showed increased expression in both LCLs from depressed subjects (supplementary figure 1g). Similar to antidepressants, n-3 PUFAs increase BDNF expression [69]. A combinatory treatment of n-3 PUFAs with escitalopram increased *BDNF* mRNA (Fig. 2l), but not proBDNF protein expression (supplementary figures 1a and 1f). Further study on the regulating *BDNF* gene and protein isoform expression might provide clarity for BDNF signaling and function in this LCLs from healthy and depressed subjects and roles of antidepressant in BDNF processing.

Alteration of the membrane microenvironment by n-3 PUFAs have been reported [70]. We could not detect altered *CAV1* expression after any treated conditions (Fig. 2b), but we did detect increased *FLOT1* expression following n-3 PUFA co-treatment with escitalopram (Fig. 2a). Both n-3 PUFAs and escitalopram redistributed flotillin-1 on non-lipid raft membrane microdomains, where it is normally less abundant (Fig. 3a and 3c). This result is consistent with previous reports showing that flotillin-1 translocated to non-lipid raft after fish oil treatment [71], suggesting effect of n-3 PUFAs on membrane protein modification and lipid raft disruption. In addition, n-3 PUFAs increased G_olf_α, but decreased G_s_α distribution on the plasma membrane of lymphoblasts (Fig. 3). Normally, G_olf_α is highly expressed in olfactory neurons where it is coupled with odorant receptors (ORs), although it is found in both the frontal cortex and lymphocytes [72]. One role for G_olf_α is to activate AC3, and this is the second most abundant adenylyl cyclase in LCLs. Furthermore, it has been reported that ORs and G_olf_α might play roles in leukocyte chemotaxis and migration [72]. It is possible that increased G_olf_α membrane distribution after n-3 PUFA treatment (Fig. 3i-3l) might play role for immune cell communications among white blood cells.

Under normal conditions, elevated cAMP suppresses/kills lymphocytes [73]. Here, we reported that n-3 PUFAs and escitalopram treatment decreased cAMP accumulation in LCLs, which may be caused by decreased G_s_α expression and altered membrane distribution. Diminished cAMP generation in LCLs (Fig. 4) might, thus, be consistent with increased cAMP in neurons or glia. Note that total cAMP accumulation might not reflect heterogeneous differences in regional cAMP within the cells. Additionally, we found up-regulation of some *ADCY* genes, as well as *BDNF,* which is regulated by cAMP response element binding protein (CREB). However, it is also possible that this was cAMP independent. Unlike neurons and glia, escitalopram and n-3 PUFAs reorganized the membrane microdomain and decreased cAMP accumulation in LCLs (Fig. 3-4). The results with the cholesterol chelator, MBCD (supplementary figure 3 and 4), may provide the answer to this conundrum. Previous studies have reported that escitalopram or MBCD treatment increased cAMP in glial cells and brain [50], yet herein, we observed that PUFAs, escitalopram and MBCD decreased cAMP in LCLs. Figure 3 shows that PUFA remove G_s_α from the membrane, leaving less G_s_α to activate AC, and decreased cAMP in LCLs (Fig. 4). These findings suggest cell type specificity for antidepressant response related to cAMP signaling, and this has been observed previously [74]. Omega-3 PUFAs have been used as anti-triglycerides and alter cholesterol levels [75], suggesting that anti- depressant-like effects of n-3 PUFAs might be due to modulation of lipid distribution and lipid metabolism. In short, n-3 PUFAs might have multiple sites of actions that can modify membrane mocrodomain and related signaling, suppress immune function, decrease cholesterol and synergize with conventional antidepressants to eventually reverse depressive symptoms.

There are some limitations for this study. Firstly, cell lines from healthy controls and depressed subjects were from different studies because the STAR*D study did not include healthy controls. Thus, we could not investigate nonspecific effects of study bias. Nonetheless, cell lines from healthy donors were picked from another study in the same cell bank, and prepared similarly to the STAR*D samples. Secondly, LCLs from the STAR*D study were collected with broad inclusion criteria [76] at various times during the course of treatment, and could alter “state-dependent” markers. Thirdly, the single concentration of Triton-X detergent extraction might not be the best option for membrane extraction because some proteins show a facultative and variable raft association during detergent extraction [77]. “Lipid –rafts” from LCLs may be different from those prepared form neurons or glia. The Triton-X 100/Triton X- 114 was chosen because it has given reliable data. Unlike sucrose gradients, Triton-X 100/Triton X-114 can be adapted for higher throughput. It is hoped that LCLs from subjects with identified depression and therapeutic response will prove useful in identifying cellular biomarkers for depression that might also aid in personalized treatment.

In summary, human lymphoblast cells might be useful models for drug discovery research for depression in order to identify the susceptibility in individuals and select the best treatment method for patients. Omega-3 PUFA alone or in combination with SSRIs exerts beneficial effects for treatment in depression. While these cells respond very differently to antidepressants than do neurons or glia, they show a reliable and predictable response that varies with both depression and antidepressant response of the subject form whom the lines were prepared.

## Acknowledgements

The authors are grateful to Athanasia Kotsouris for help with ELISA. This project was supported by NIH Grant R41MH113398 and R01AT009169 and VA grant BX001149. MMR is a VA Career Research Scientist BX004475. Phatcharee Chukaew was also supported by a Science Achievement Scholarship of Thailand (SAST).

