## Supplementary figures and images for "Potential depression and antidepressant-response biomarkers in human lymphoblast cell lines from treatment-responsive and treatment-resistant subjects: roles of SSRIs and omega-3 polyunsaturated fatty acids"

### Fig S1

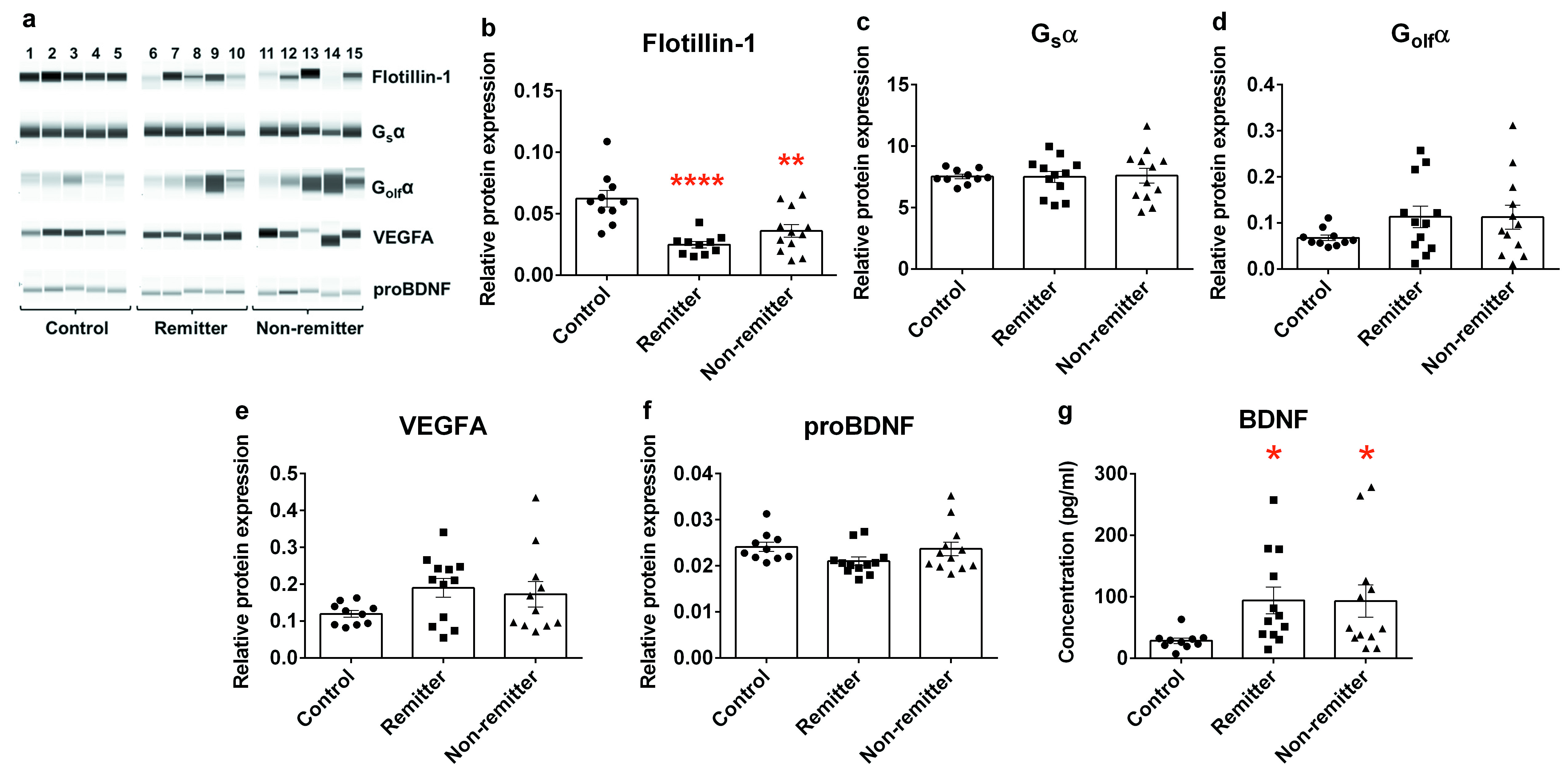

### Fig S2

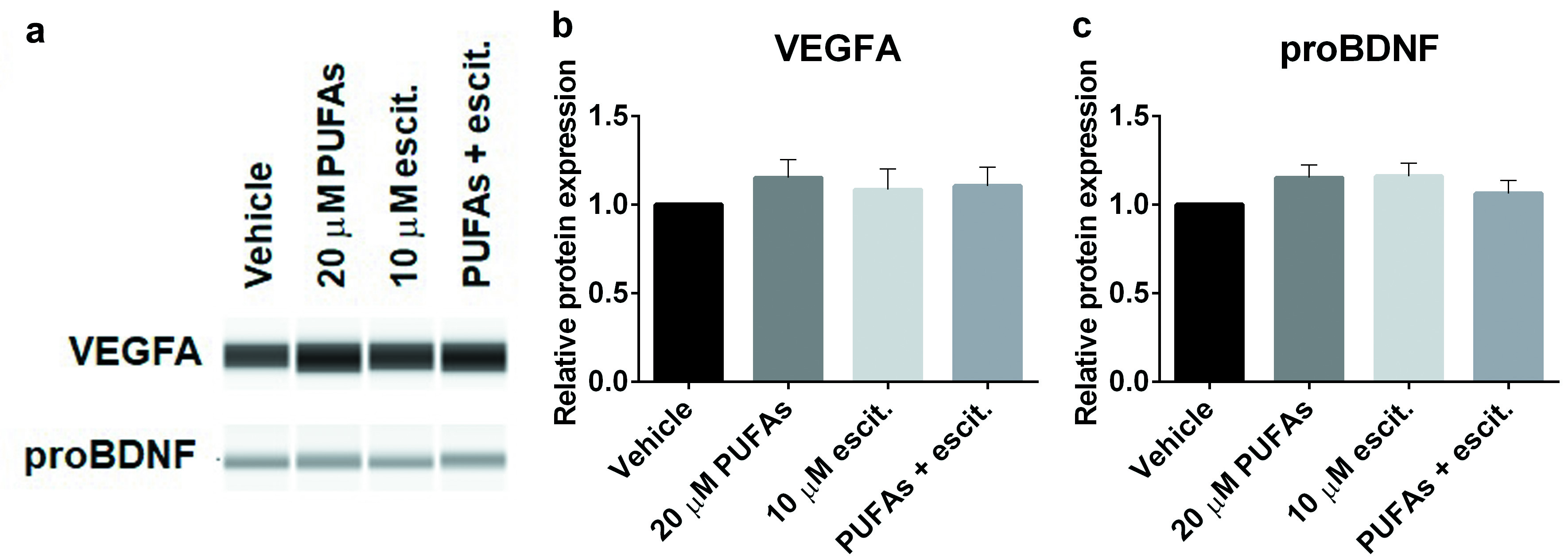

### Fig S3

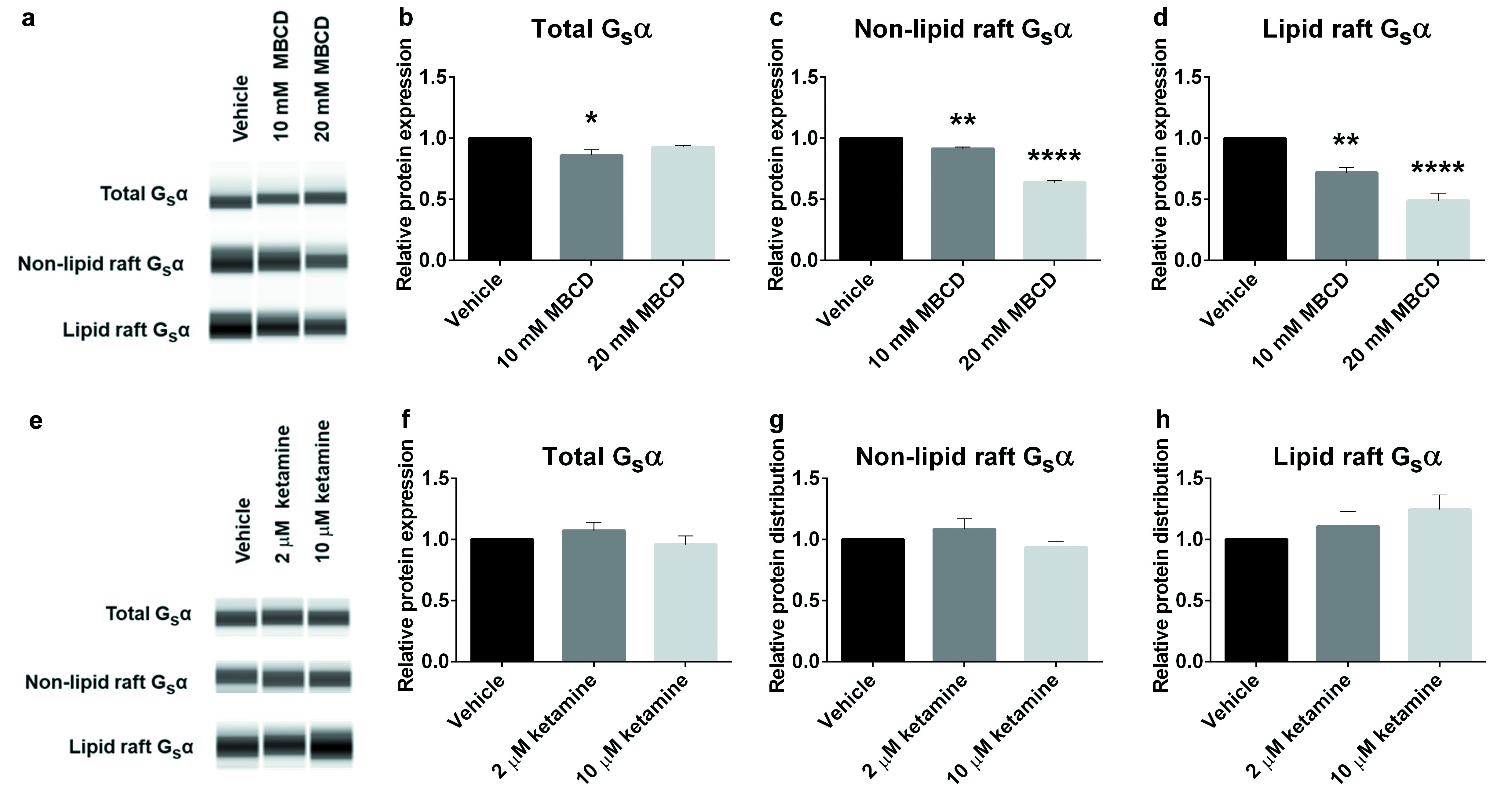

### Fig S4

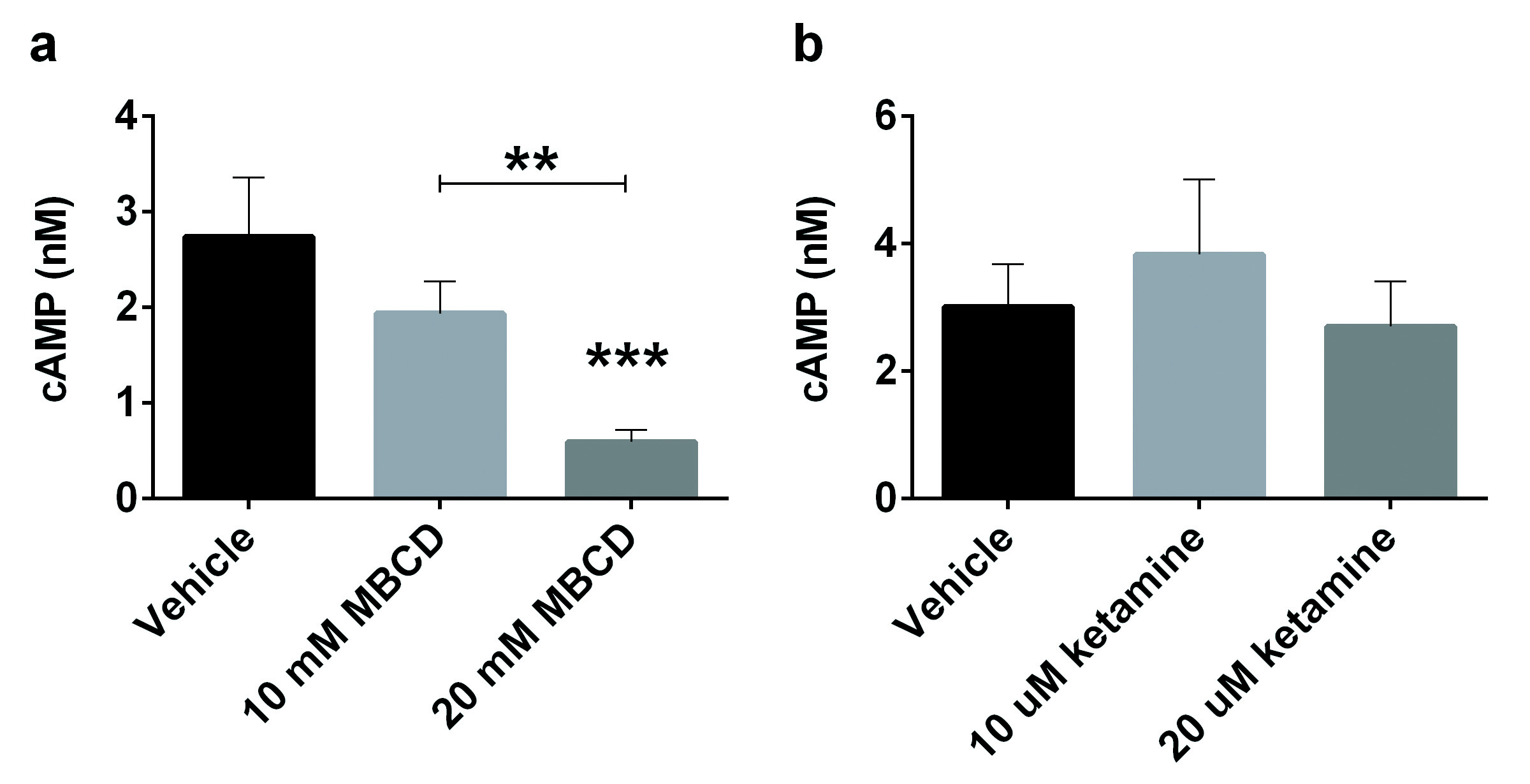
