## Supplemental methods and legends for "Potential depression and antidepressant-response biomarkers in human lymphoblast cell lines from treatment-responsive and treatment-resistant subjects: roles of SSRIs and omega-3 polyunsaturated fatty acids"

**Supplementary data**

**Supplementary figure 1** Protein expression in LCLs from healthy controls and depressed subjects who remitted and did not remit to antidepressant treatment. Lysates were collected using RIPA lysis buffer. Protein concentrations of each sample were equalized before loading the same amount into a Protein Simple Wes cassette for capillary electrophoresis and immunodetection. **a** The expression level of protein represented by western blotting lanes for each individual sample and mean ± SEM including **b** Flotillin-1, **c** G_s_α, **d** G_olf_α, **e** VEGFA and **f** proBDNF. **g** total BDNF levels were measured with BDNF ELISA and depicted as mean ± SEM. Adjusted *p* value from 3 comparisons among groups of cell lines, **p*<0.05, ***p*<0.01, *****p*<0.0001.

**Supplementary figure 2** PUFAs and escitalopram had no effect on VEGFA and proBDNF protein expression in LCLs. Lysates were collected from LCLs that were treated with vehicle, 20 μM PUFAs, 10 μM escitalopram or 20 μM PUFAs + 10 μM escitalopram for 72 hours. Protein concentrations of each sample were equalized before immunodetection. The experiment was done in 9 cell lines derived from depressed subjects. **a** The expression level of protein from one cell line is depicted. Data from 9 cell lines are quantified as and represented as mean ± SEM for **b** VEGFA and **c** proBDNF.

**Supplementary figure 3** Methyl-β-cyclodextrin (MBCD), but not ketamine, decreases membrane distribution of Gs_α_ in LCLs from depressed subjects. LCLs were treated with vehicle, 10 or 20 mM MBCD for 30 minutes, and 2 or 10 μM ketamine for 60 minutes. Protein concentrations of each sample were equalized before capillary electrophoresis and immunodetection **a-d** MBCD (n=4) and **e-h** ketamine (n=13) treatment. Values indicate as bars of mean ± SEM. Adjusted *p* value of 3 comparisons between each pair of treated conditions, **p*<0.05, ***p*<0.01, *****p*<0.0001.

**Supplementary figure 4** Methyl-β-cyclodextrin (MBCD), but not ketamine, decreases cAMP accumulation in LCLs derived from depressed subjects. LCLs were treated with vehicle, 10 or 20 mM MBCD for 30 minutes, 10 or 20 μM ketamine for 60 minutes. Alpha screen was used to measure cAMP accumulation in cells. The experiment was done in triplicates for 3 cell lines. **a** MBCD but not **b** ketamine decreased cAMP accumulation in LCLs. Values shown as mean ± SEM. Adjusted *p* value of 3 comparisons between each pair of treated conditions, ***p*<0.01, ****p*<0.001.

**Supplementary Table 1** Primers for target genes used in this study

| **Gene** | **Encode protein** | **Forward primer ( 5’🡪3’)** | **Reverse primer ( 5’🡪3’)** |
| --- | --- | --- | --- |
| ***FLOT1*** | Flotillin-1 | AAGCTGCCCCAGGTGGCAGAGG | TGTTCTCAAAGGCTTGTGATTCACC |
| ***CAV1*** | Caveolin-1 | CTGACATTCCCTGATGACGCTG | AGATGTGCAGGAAAGAGAGAATG |
| ***GNAS*** | G_s_ alpha subunit, G_s_α | TGCTGGAGAATCTGGTAAAAG | CCTTGGCATGCTCATAGAATTC |
| ***GNAL*** | G_olf_ alpha subunit, G_olf_α | CAGCGTCAGCTTGGTTGACTA | GTAATGTTTGCCGTCACCGGT |
| ***ADCY1*** | Adenylyl cyclase 1, AC1 | GCGCAGCTACGAGCCGATTG | AGGAAGTGCTGGGCGACGTG |
| ***ADCY3*** | Adenylyl cyclase 3, AC3 | CGGTGGAGAAGGAGAAGCAGAGTGG | CCTCCGTCTCCATCCCTGCCGTTGC |
| ***ADCY6*** | Adenylyl cyclase 6, AC6 | GTTGCCTGTGCCCTGTTGGTCTTCT | GATGCCAACTGCGGTGCTATGTGCC |
| ***ADCY7*** | Adenylyl cyclase 7, AC7 | ACTGGGTGTGTCCTTCGGGCTGGTG | CTCTGGAACTTGGAAAGGCATCAGG |
| ***ADCY9*** | Adenylyl cyclase 9, AC9 | TGGGAAAGTTATTGAACGGCTG | CTGACATTCCCTGATGACGCTG |
| ***VEGFA*** | Vascular endothelial growth factor A, VEGFA | CTGTACCTCCACCATGCCAAG | GGTACTCCTGGAAGATGTCCACC |
| ***IL6*** | Interleukin 6, IL6 | GGTACATCCTCGACGGCATCT | GTGCCTCTTTGCTGCTTTCAC |
| ***BDNF*** | Brain-derived neurotrophic factor, BDNF | GATGAGGACCAGAAGGTTCG | TCCAGCAGAAAGAGCAGAGG |
| ***ACBT*** | Beta actin | GGGAGAAGCATTAGGAGGG | CAAGGAAAGCAAGGTGGG |

**Supplementary table 2** Demographic information of human lymphoblast cell lines from healthy control and depressed subjects who remitted or failed to remit to sequenced treatment

| **Group** | **Sample ID** | **Sex** | **Age** | **Race** |
| --- | --- | --- | --- | --- |
| **Control** | 09C81954 | Male | 55 | White |
|  | 09C89787 | Male | 38 | White |
|  | 09C89437 | Male | 42 | Mixed |
|  | 09C89740 | Male | 46 | White |
|  | 09C90778 | Male | 22 | Black |
|  | 09C81021 | Female | 55 | White |
|  | 09C81052 | Female | 32 | Mixed |
|  | 09C89398 | Female | 26 | White |
|  | 09C90516 | Female | 22 | White |
|  | 09C89961 | Female | 28 | Black |
| **Remitter** | 04C26645 | Male | 22 | White |
|  | 03C14548 | Male | 33 | White |
|  | 04C24599 | Male | 63 | White |
|  | 03C19754 | Male | 37 | Mixed |
|  | 03C20744 | Male | 43 | White |
|  | 03C19634 | Female | 36 | Black |
|  | 03C20212 | Female | 55 | Black |
|  | 04C26525 | Female | 42 | Black |
|  | 04C27584 | Female | 36 | White |
|  | 03C19958 | Female | 53 | White |
|  | 03C21151 | Female | 28 | Mixed |
|  | 04C33393 | Female | 34 | White |
| **Non-remitter** | 03C18967 | Male | 47 | White |
|  | 04C27349 | Male | 39 | White |
|  | 03C16750 | Male | 28 | White |
|  | 03C21487 | Male | 46 | Mixed |
|  | 04C24601 | Male | 57 | White |
|  | 04C28940 | Male | 51 | White |
|  | 03C21947 | Female | 62 | Mixed |
|  | 03C16752 | Female | 32 | White |
|  | 03C17862 | Female | 54 | White |
|  | 04C25339 | Female | 53 | White |
|  | 03C20107 | Female | 54 | Black |
|  | 03C22174 | Female | 49 | Black |

**Supplemental table 3** Clinical characteristics of MDD lymphoblast cell line donors

| **Group** | **Sample ID** | **Episode number^a^** | **Drug usage^b^** | **Personal disorders^c^** | **STAR*D level^d^** | **HRS score** | | |
| --- | --- | --- | --- | --- | --- | --- | --- | --- |
|  |  |  |  |  |  | **Enrollment** | **Citalopram treatment (≤ 14 weeks)** | **Antidepressant treatments (≤ 56 weeks)** |
| **Remitter** | 04C26645 | - | 1 | 2 | 1 | 17 | 0 | 0 |
|  | 03C14548 | 1 | 2 | 1 | 2 | 15 | 11 | 5 |
|  | 04C24599 | 3 | 2 | 2 | 2 | 15 | 8 | 4 |
|  | 03C19754 | - | 2 | 2 | 3 | 17 | 11 | 5 |
|  | 03C20744 | 2 | 2 | 2 | 4 | 18 | 17 | 5 |
|  | 03C19634 | 2 | 1 | 1 | 1 | 16 | 2 | 2 |
|  | 03C20212 | - | 2 | 2 | 1 | 16 | 3 | 3 |
|  | 04C26525 | 2 | 1 | 1 | 2 | 17 | 6 | 1 |
|  | 04C27584 | 1 | 1 | 2 | 2 | 16 | 7 | 6 |
|  | 03C19958 | 1 | 1 | 1 | 4 | 18 | 7 | 6 |
|  | 03C21151 | 3 | 2 | 2 | 3 | 17 | 20 | 7 |
|  | 04C33393 | - | 2 | 2 | 3 | 17 | 10 | 2 |
| **Non-remitter** | 03C18967 | 51 | 1 | 2 | 1 | 15 | 10 | 10 |
|  | 04C27349 | 1 | 1 | 1 | 2 | 17 | 10 | 10 |
|  | 03C16750 | 4 | 2 | 1 | 3 | 18 | 8 | 8 |
|  | 03C21487 | - | 1 | 1 | 2 | 14 | 25 | 19 |
|  | 04C24601 | 10 | 2 | 2 | 3 | 18 | 14 | 16 |
|  | 04C28940 | - | 1 | 1 | 4 | 17 | 15 | 18 |
|  | 03C21947 | 4 | 2 | 1 | 4 | 17 | 20 | 13 |
|  | 03C16752 | 2 | 2 | 2 | 3 | 18 | 9 | 9 |
|  | 03C17862 | 1 | 1 | 2 | 4 | 16 | 16 | 10 |
|  | 04C25339 | 1 | 2 | 2 | 1 | 15 | 15 | 15 |
|  | 03C20107 | 2 | 1 | 1 | 2 | 18 | 24 | 27 |
|  | 03C22174 | 6 | 1 | 1 | 4 | 16 | 23 | 28 |

- = not applicable

a = number of depressed episodes at enrollment 51??

b = experience of drug usage (alcohol, amphetamine, cocaine, cannabis or opioid) from DSM-4 AXIS II diagnoses (1 = no, 2= yes)

c = personal disorders from DSM-4 AXIS II diagnoses (1 = no, 2= yes) personality??

d = STAR*D level refers to assigned drug to participants followed STAR*D study protocol. All subjects initially received citalopram up to 14 weeks. If a participant did not become symptom‐free or could not tolerate side effects from the citalopram, he/she was

encouraged to progress to the next level.
